# Genomic Tokenizer: Toward a biology-driven tokenization in transformer models for DNA sequences

**DOI:** 10.1101/2025.04.02.646836

**Authors:** Bell Raj Eapen

**Affiliations:** University of Illinois Springfield, Springfield, IL 62703

**Keywords:** Tokenizer, Transformer, DNA

## Abstract

**Summary:** Transformer models are revolutionizing sequence analysis across various domains, from natural language processing to genomics. These models rely on tokenizers to split input sequences into manageable chunks — a straightforward task in natural language but more challenging for long DNA sequences that lack distinct “words.” Most biological tokenizers are data-driven and do not align with the “central dogma of molecular biology”: DNA is transcribed into RNA, which is then translated into proteins, with each three-letter codon specifying a particular amino acid, some of which are synonymous for the amino acids they represent. Start codons signal the beginning of protein synthesis, while stop codons signal its termination. The Genomic Tokenizer (GT) incorporates this biological process flow into a standard tokenizer interface within the HuggingFace transformer package. GT can be used to pre-train foundational transformer models on DNA sequences. We compare the performance of GT with two alternate tokenization strategies and discuss its potential applications.

**Availability and implementation:** The source code of GT is available from https://github.com/dermatologist/genomic-tokenizer under the MPL-2.0 license. It can be installed from Python Package Index (PyPI) and used as a tokenizer in transformer model training pipelines.

## 1 Introduction

The success and popularity of the use of transformer-based large language models (LLMs) in various domains have led to the exploration of the transformer architecture and attention mechanism in genomic sequence analysis. However, genomic sequences, whether DNA, RNA, or protein, vary from a sequence of words in human languages in many ways. Genomic sequences, unlike human language, lack distinct words, are highly repetitive, and have a complex structure that includes genes and regulatory elements. Despite these differences, models leveraging transformer architectures can still be beneficial for genomic sequence analysis due to their ability to capture complex patterns and dependencies within the sequence data. The attention mechanism [1] allows models to focus on relevant parts of genomic sequences, which is crucial when dealing with large-scale biological datasets where specific regions may hold critical information regarding gene expression or mutations associated with diseases. However, genomic sequences can be extremely long, making it computationally expensive and memory-intensive for standard transformers to process them in their entirety.

The initial step in processing any sequence with a transformer architecture is to divide the sequence into small, manageable chunks known as tokens. Unlike natural language, which can be tokenized based on words, tokenizing DNA sequences presents unique challenges as meaningful units are not clearly defined. A simple character tokenization method is to consider each nucleotide (A, G, C, and T) as a single token. A, G, C, and T stand for the nucleotides: adenine, guanine, cytosine, and thymine, respectively, in DNA sequences. This approach makes the transformer models attend to every nucleotide and may fail to capture higher-order information, especially in long sequences.

An alternative approach, k-mer tokenization, breaks the sequence into overlapping or non-overlapping segments of a predefined length k. While k-mer tokenization captures more contextual information than single-nucleotide tokenization, the optimal k value is often context-dependent and requires careful consideration. Several advanced data-driven approaches, such as byte-pair encoding (BPE) [2], have been proposed, with varying degrees of success [3]. If annotations are available, high-level features such as gene regulatory elements can be used as tokens [4]. Data-driven methods can be trained on co-occurrence patterns within a large corpus of DNA data [5]. However, most techniques are resource-intensive and not widely generalizable. Developing algorithms for efficient tokenization of DNA remains an active area of research, and the choice of tokenization strategy depends heavily on the specific downstream application.

According to the central dogma of molecular biology [6], DNA is first transcribed into RNA (specifically mRNA), which then serves as a template for protein synthesis during translation. Codons are sequences of three nucleotides (k=3) that correspond to a specific amino acid or a stop signal during protein synthesis. Transcription begins with a start codon (typically AUG) and continues through several codons (reading frames), each representing specific amino acids, till one of the three stop codons is encountered. Some of the codons are synonymous as they code for the same amino acid. Exons are the gene’s coding regions that are expressed, whereas introns are non-coding regions that are excised during RNA processing. A mutation is a change in the DNA sequence that can lead to changes in protein structure and function. Addition or deletion of even a single nucleotide can shift reading frames, leading to extensive changes in the resulting protein. Single nucleotide polymorphisms (SNPs) are variations at a single position in the DNA sequence among individuals. These variations can influence gene expression and contribute to diverse phenotypic traits and diseases. In summary, the biology of DNA sequences is fundamentally different from the structure of word sequences.

Genomic Tokenizer (GT) incorporates start codons, synonymous codons, and stop codons into a tokenizer interface of the HuggingFace transformer package [7], giving it the ability to handle shifts in reading frames caused by nucleotide additions or deletions within DNA sequences. By doing so, GT ensures that biological nuances inherent in genetic variations are preserved during tokenization, potentially enhancing their performance on tasks related to phenotypic predictions.

## 2 Materials and methods

Hugging Face Transformers is an open-source Python library that provides tools for manipulating transformer models. It includes tokenizers implementing methods for splitting a string into chunks and encoding them into numerical representations (token IDs) and vice versa for input into a transformer model [8]. The core functionality takes a string or a list of strings as input and returns a dictionary containing the token IDs (input_ids) and the attention mask used to differentiate between actual data and padding. The *PreTrainedTokenizer* is a general-purpose Python base class [9] from which specific tokenizer classes such as *BertTokenizer* and *GPT2Tokenizer* inherit from implementing the respective tokenization strategies. Each tokenizer uses a vocabulary – a mapping between tokens and their corresponding IDs – specific to the tokenization strategy. Additionally, tokenizers manage special tokens like CLS, SEP, PAD, UNK, and MASK, each serving specific roles in transformer models. These roles include marking the start of a sequence, separating segments, padding shorter sequences, representing unknown tokens, and masking tokens for prediction tasks respectively.

The GenomicTokenizer class extends the PreTrainedTokenizer class and implements the required interfaces. The vocabulary includes all possible codons, but synonymous codons that code for the same amino acids are assigned the same IDs, effectively reducing the vocabulary size and improve the efficiency of the tokenizer. The IDs one to six are reserved for special tokens. “ATG” is assigned as the start codon and “TAA”, “TAG” and “TGA” as the stop codons. These are customizable for use in the context of certain prokaryotic genomes where they differ. Start codon is treated as the **BOS** token and the end codons as **SEP** tokens. Tokenization begins at the start codon if one is identified within the sequence. If no start codon is found, tokenization defaults to the beginning of the sequence. If an end codon is found, all subsequent codons are marked as **UNK** tokens until another start codon is encountered. **UNK** tokens at the end are trimmed off so that padding can be applied as required. This process ensures that only the presumed coding sequence, delineated by start and stop codons, is processed and assigned meaningful tokens. The **UNK** tokens are attended to, but do not contribute towards loss calculation. The coding of introns as **UNK** tokens can be turned off to reduce the token count further. The tokenization algorithm is summarized in Table 1.

**Table 1:**
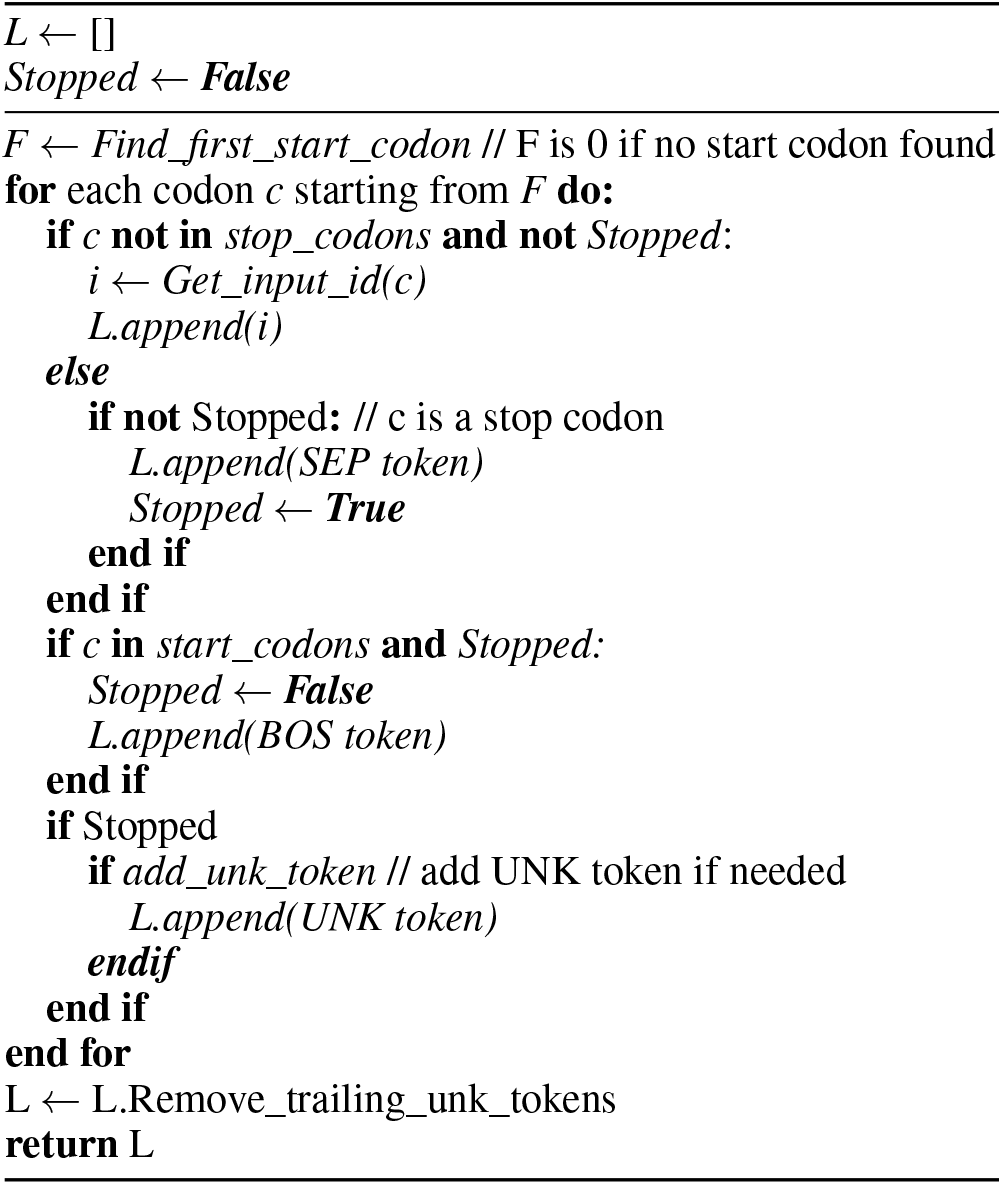
The tokenization algorithm.

## 3 Result

The complexity of genomic data, and variability of dataset and downstream tasks makes comparison of tokenizers difficult. We present a preliminary comparison of GT with two other tokenizers; HyenaDNA’s character tokenizer [10] (henceforth CT) with a vocabulary size of 4 and DNABERT-2’s [5, 11] data driven adaptation of SentencePiece [12] (henceforth BPE) with a vocabulary size of 2048. We used GV-Rep; a pipeline for creating clinician-verified genetic variant sequences of specified lengths from reference genome [13]. Using GV-Rep we generated a subset of sequences linked to lung cancer as positive samples and an equal number of sequences representing non-lung cancer conditions as negative samples from ClinVar (a public archive of reported variants associated with diseases) [14] with sequence lengths; 512, 1024, 2048 and 4096 for the comparison of tokenizers. Using this generated dataset, we trained a simplified BERT architecture for sequence classification from scratch. We used the following hyperparameters throughout; a learning rate of 3e-4, and a batch size of 12 with 3 hidden layers and 3 attention heads each. The hidden layer dimension was 192 and the optimizer AdamW was used to train the model for one epoch.

We compared three classification evaluation metrics: Accuracy (ACC), Area Under the Curve (AUC) and Matthew’s Correlation Coefficient (MCC) for CT, BPE and GT tokenizers using the BERT model and dataset described above. Accuracy (ACC) measures the proportion of true positive and true negative predictions among the total number of cases, reflecting the overall correctness of a model. The Area Under the Curve (AUC) of the Receiver Operating Characteristic (ROC) curve quantifies the ability of a model to distinguish between positive and negative classes, with higher values indicating better discrimination. Matthew’s Correlation Coefficient (MCC) measures the quality of binary classifications, especially for unbalanced datasets. It ranges from -1 to +1, where +1 means perfect prediction, 0 means random guessing, and -1 means total disagreement between predictions and actual outcomes.

Data-driven BPE performed best in this task, though direct comparison of tokenizers is unjust due to the differences in vocabulary size, which in turn leads to minor changes in the BERT architecture used for comparison. A larger vocabulary size typically results in an increased number of trainable parameters, as each distinct word in the vocabulary needs its own embedding vector. The CT’s performance steadily decreased with increasing sequence length, exhibiting a clear sensitivity to the length of the input sequence. This suggests that the fixed-length representation inherent in character-level tokenization struggles to capture the long-range dependencies crucial for understanding longer sequences. In contrast, GT demonstrated greater robustness to variations in sequence length (see Figure 1 B-D).

**Figure 1.**
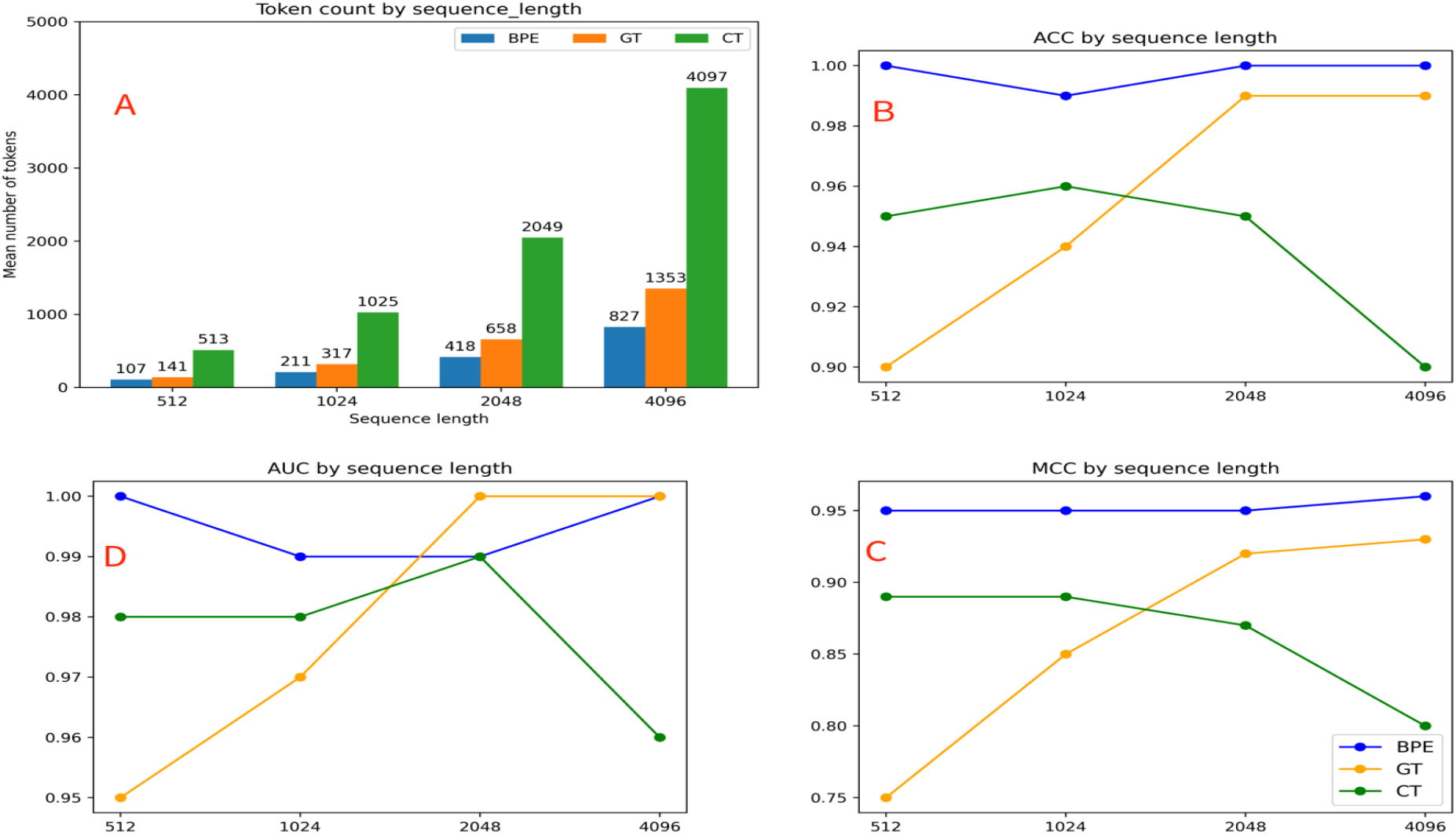
Comparison showing A: Number of tokens generated and B,C,D: Accuracy (ACC), Matthews Correlation Coefficient (MCC) and Area Under Curve (AUC) for BPE, GT and CT tokenizers against the sequence length.

Additionally, we have compared the average token count for each tokenizer. CT treats each character as a separate token, while BPE tokenization merges frequent character pairs iteratively, creating tokens that represent more than one character. GT adopts a 3-mer tokenization that accounts for synonymous codons and exons. As expected, the average token count of GT was lower compared to the character tokenizer but higher than BPE (see Figure 1A). Furthermore, a simplified BERT model for masked language modeling (MLM) demonstrated the highest accuracy for CT, the lowest for BPE, and intermediate results for GT, consistent with the vocabulary size for each tokenizer.

## 4 Discussion

The focus of data driven tokenizers is on finding repetitive elements in DNA sequences reducing the number of tokens to process. However, this leads to an increase in the size of the dictionary making them computationally resource intensive compared to non-data driven tokenizers such as CT or k-mer tokenizers. Furthermore, despite the apparent simplicity of having only four nucleotides as alphabets, the information that DNA encodes is highly variable across organisms, chromosomes and genomic regions, many of which are still unknown. GT’s biology-driven algorithm has the potential to maintain compact vocabulary, which significantly enhances computational efficiency. Additionally, it may capture long token dependencies crucial for modeling the complex and intricate relationships within DNA sequences. This needs to be confirmed by using GT for foundational model training.

Overlapping k-mer tokenization leads to considerable redundancy and information leakage in masked language modelling (MLM) when adjacent tokens are not masked. As the vocabulary size increases, the search space in MLM increases along with the computational complexity. GT does not have either of these potential drawbacks.

Specific mutations or genetic variations in the DNA sequence can affect an organism’s phenotype (observable characteristics). These alterations can range from single nucleotide changes (point mutations) to large-scale chromosomal rearrangements. Even single nucleotide polymorphisms (SNPs) and mutations can significantly influence biological properties [15]. Point mutations can be substitution, insertion or deletion of the nucleotide. If the substitution is “synonymous”, it will not alter the amino acid sequence of the resulting protein due to the redundancy of the genetic code. Missense change resulting in a different amino acid being incorporated into the protein can have varying effects, from no noticeable change to complete loss of function of the protein. “Nonsense” mutation creating a premature stop codon leads to a truncated protein. Insertion or deletion of one or more nucleotides can cause a frameshift altering the amino acid sequence downstream leading to a completely different protein. Understanding specific mutations and genetic variations is crucial for diagnosing and treating genetic diseases, developing personalized medicine, and understanding evolutionary processes. GT can capture these biological variations during model training. Additionally, start and stop codons can be customized to accommodate variations in mitochondrial genomes and certain prokaryotic organisms. Furthermore, customizable intron encoding enhances its utility for various tasks.

No single tokenization strategy is optimal for all datasets [3] and the optimal choice depends on several factors such as the dataset, task, and the computational resources available. GT may be useful in existing architectures such as the convolutional long-context model of HyenaDNA [10]. Ultimately, a thorough comparative analysis across various tokenization methods, vocabulary sizes, and base models is crucial for determining the most effective approach for a given application. We offer GT to the open-source community, encouraging exploration with diverse datasets and tasks.

## 5 Acknowledgements

We gratefully acknowledge the infrastructural support provided by Orion Lab at the University of Illinois Springfield (UIS).

## Notes

### Competing Interest Statement

The authors have declared no competing interest.

